# The larval *Drosophila* mushroom body balances lateralized sensing and interhemispheric integration

**DOI:** 10.1101/2025.03.02.641007

**Authors:** David M. Zimmerman, Benjamin L. de Bivort, Aravinthan D. T. Samuel

## Abstract

All animals with bilateral symmetry must integrate the sensory input from the left and right sides of their bodies to make coherent perceptual decisions. In the *Drosophila* larva, olfactory receptor neurons project largely ipsilaterally, providing a tractable system for asking where and how interhemispheric integration arises downstream. We combined volumetric calcium imaging with unilateral sensory perturbations, connectomic analysis, and optogenetic manipulations to trace the propagation of left–right olfactory information across successive layers of the olfactory system. This approach implicates the mushroom body (MB) as a key substrate for interhemispheric integration of odor representations. Kenyon cell (KC) odor responses were almost entirely ipsilateral, indicating minimal functional coupling between the two MBs at the input level. In contrast, modulatory neurons (MBINs) exhibited highly symmetric responses to unilateral stimulation, suggesting that reinforcement signals are broadly shared across hemispheres. Nevertheless, odor responses in some MB output neurons (MBONs), up to 5 synapses downstream from the sensory periphery, preserve information about stimulus laterality. Moreover, we show that asymmetric activation of these MBONs can modulate the animal’s turning behavior in a side-biased manner. Finally, we provide direct evidence that larvae can exploit instantaneous spatial comparisons for navigation in certain sensory contexts. These findings suggest that the deeply lateralized architecture of the larval olfactory system balances the need for interhemispheric integration with the advantages of parallel sensory processing.

## 1 Introduction

Any animal with bilateral symmetry must combine the two parallel streams of sensory input originating from the two sides of its body into a single unified representation of the world. Such interhemispheric integration (IHI) is a necessary prerequisite for transforming spatially and temporally disparate sensory signals into coherent motor commands. Neurological conditions in which IHI is compromised invariably lead to severely impaired perceptual decision making and disordered sensorimotor coordination (*1*). Even so, it is often advantageous for animals to process bilateral sensory signals in such a way that their spatial structure can be exploited. For instance, many animal species use interocular disparities to perceive depth (*2*), interaural timing differences to localize sounds (*3*), and internasal odor gradients to navigate in chemical landscapes (*4*). The ability to selectively integrate or compare sensory information across hemispheres allows animals to sharpen their sensitivity to stimuli and adapt their behavioral responses to dynamic environments. However, it is poorly understood how bilaterian nervous systems implement these two different types of sensory processing in a balanced and mutually compatible way.

Recent work in rodent and primate models has begun to explore the circuit mechanisms underlying flexible IHI of somatosensory (*5, 6*), auditory (*7, 8*), and olfactory maps (*9* –*11*). The case of mammalian olfaction has attracted particular interest due to the apparently ipsilateral nature of the afferents to cortex combined with the lack of topographic organization in piriform odor representations. These neuroanatomical features are difficult to reconcile with demonstrations of interhemispheric transfer of learned olfactory associations and reports that individual cortical neurons exhibit similar tuning to both ipsilateral and contralateral stimuli (*9*). Several alternative hypotheses have been suggested, including indirect excitatory coupling between cognate glomeruli in the left and right olfactory bulbs and sculpting of interhemispheric connectivity at the cortical level by experience-dependent plasticity (*10, 11*). Unfortunately, the sheer size and complexity of the mammalian brain, the lack of a detailed wiring diagram for the mouse olfactory system, as well as the practical limitations of currently available neurogenetic tools, have hindered efforts to test these hypotheses in a precise and systematic way.

The olfactory system of the *Drosophila* larva is strikingly similar in its basic organization to that of most vertebrates. In *Drosophila* as in mammals, all ORNs expressing a given olfactory receptor (OR) type converge onto a corresponding glomerulus in the first olfactory neuropil, the antennal lobe (AL), which is the insect counterpart of the mammalian olfactory bulb. However, whereas ORNs in the antennae of the adult fly project bilaterally to both the left and right ALs (*12*), ORNs in the left and right dorsal organs (DOs) of the larva project almost exclusively to the ipsilateral AL (Fig. 1A), as in mammals. The larval olfactory system also has the remarkable property that each of the 21 ORN types is instantiated by a single left/right pair of cells. As a result, the input to any given glomerulus can be completely controlled by manipulating a single sensory neuron.

**Figure 1:**
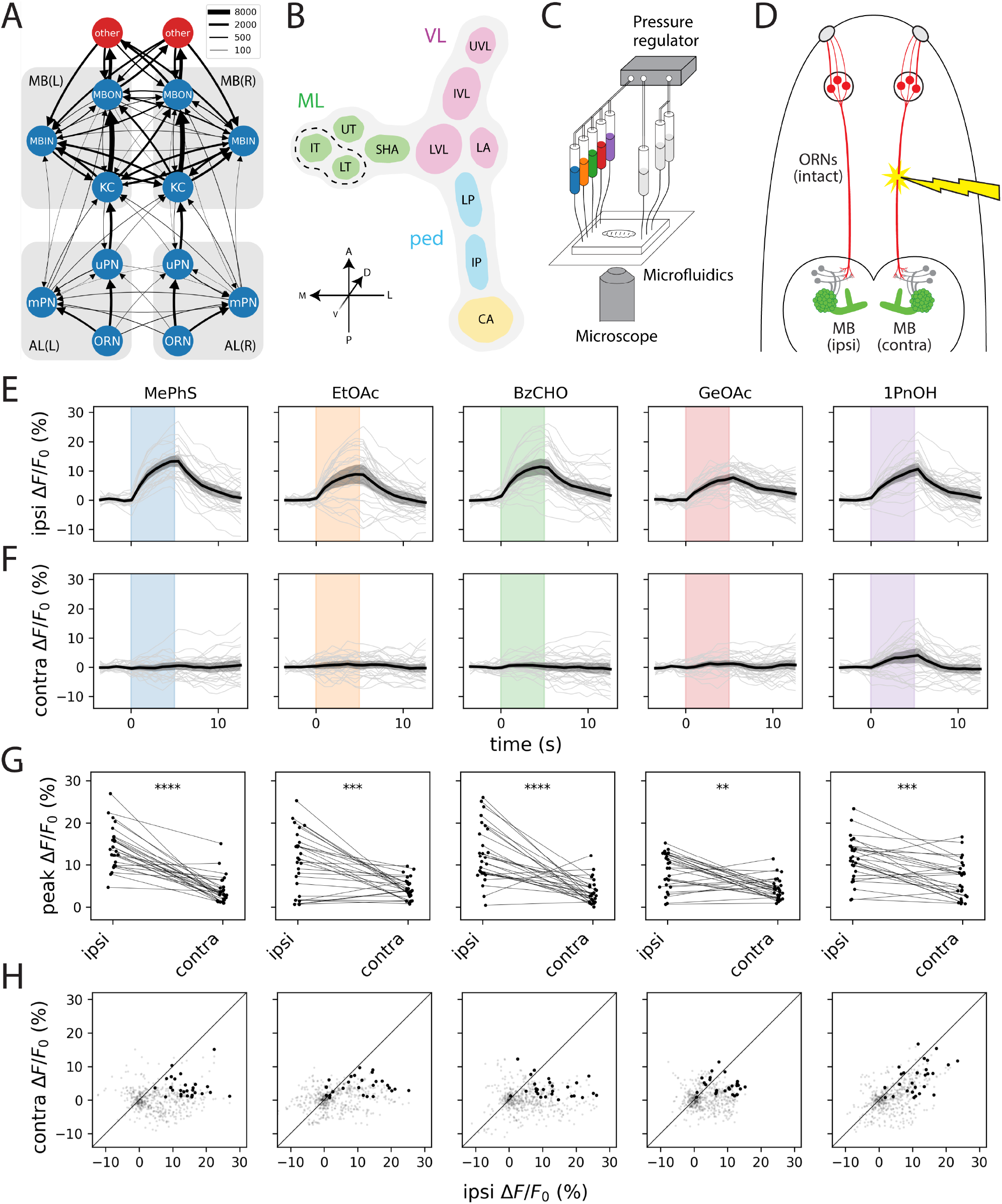
The larval MB is the first major site of olfactory IHI. A. Summarized wiring diagram of the L1 larval olfactory system (*13, 19*). Each node agglomerates all cells of the same class (ORNs, uPNs, mPNs, KCs, MBINs, MBONs) in a given hemisphere. Edge width is proportional to the square root of the number of synapses between all cells in a given pair of cell types. The red nodes labeled “other” agglomerate all cells downstream of the MB in the left and right hemispheres, respectively. See also Fig. S1. B. Simplified anatomical representation of the larval MB, consisting of the calyx (CA), pedunculus (ped), vertical lobe (VL), and medial lobe (ML), of which the latter three are further subdivided into functional compartments: intermediate pedunculus (IP), lower pedunculus (LP), lateral appendix (LA), lower vertical lobe (LVL), intermediate vertical lobe (IVL), upper vertical lobe (UVL), shaft (SHA), upper toe (UT), intermediate toe (IL), and lower toe (LT). For the purposes of this study, we group the IT and LT together (dashed lines), as we cannot reliably separate them in our pan-MB imaging experiments. C. Schematic of the experimental setup for microfluidic odor delivery during imaging. A pressure regulator drives odorant solutions through a microfluidic chip while odor-evoked calcium responses from an immobilized larva are recorded using a spinning-disk confocal microscope. Stimulus timing is controlled by a bank of computer-operated solenoid valves coupled to the odor syringes. D. Experimental strategy for synthesizing lateralized odor stimuli by laser ablation of the antennal nerve (red). The KCs of the larval MB are shown in green to represent transgenic expression of GCaMP for calcium imaging. The side ablated (left or right) was randomized across animals; ipsilateral and contralateral data are pooled across left/right experiments. E–H. KCs exhibit consistent ipsilateral and negligible contralateral odor responses; see also Fig. S2. Plots depict ipsilateral (E) and contralateral (F) odor-evoked cytosolic GCaMP6m transients and peak ipsilateral vs. contralateral response amplitudes (G). In E and F, dark lines represent grand averages over *n* = 9 animals and 3 stimulus repetitions per animal; gray shaded regions are bootstrapped 95% CIs. Colored vertical bars represent 5 s stimulus periods. F. Scatter plots of ipsilateral and contralateral KC activity in the 15 s following stimulus onset. Bold points represent peak responses. Diagonal lines represent equality between ipsilateral and contralateral response amplitudes. Each column shows responses to a different odor. All odors were used at 10^*−*4^ (v/v) dilution. Odor abbreviations: MePhS, methyl phenyl sulfide (blue); EtOAc, ethyl acetate (orange); BzCHO, benzaldehyde (green); GeOAc, geranyl acetate (red); 1PnOH, 1-pentanol (violet). Statistics: **, *p*_adj_ < 0.01; ***, *p*_adj_ < 0.001; ****, *p*_adj_ < 0.0001 (paired Student’s *t*-test with Holm’s correction).

In the AL, a population of 21 uniglomerular projection neurons (uPNs) innervate the 21 olfactory glomeruli in a one-to-one manner and project ipsilaterally (with one exception) to two higher brain areas: the mushroom body (MB) and lateral horn (LH). (Only one pair of uPNs, corresponding to the Or35a input channel, receive input from both the ipsi- and contralateral DOs and send output to downstream areas in both the ipsi- and contralateral hemispheres.) The complete wiring diagram of the first-instar (L1) larval AL also revealed the existence of 14 multiglomerular projection neurons (mPNs), which form synapses with multiple ORN types and project to a diverse set of higher brain areas (*13*). Notably, 4 mPNs per AL receive some degree of bilateral input, and 3 of these also produce bilateral output. In total, however, ORNs form 18x more synapses with ipsilateral PNs than with contralateral PNs (*13*). Thus, the larval AL circuitry is overwhelmingly unilateral in character.

In contrast, the wiring diagram of the larval MB suggests a much greater degree of interhemi-spheric crosstalk (Fig. 1A). The L1 larval MB contains approximately 100 intrinsic neurons, called Kenyon cells (KCs), each of which receives input from a random subset of 1–6 PNs (*14*). This population of KCs then converges onto a multilayered recurrent network of 24 MB output neurons (MBONs), which send output to many regions of the dorsal brain. The MBON dendrites arborize in spatially disjoint compartments (Fig. 1B) collectively tiling the KC axons in such a way that every KC forms en passant synapses with most MBONs. Each compartment is also innervated by 1–3 MB input neurons (MBINs), capable of heterosynaptically modulating the strength of the corresponding KC-MBON connections in response to positive or negative reinforcement according to a compartment-specific learning rule (*15*).

Previous studies have implicated multilayered connectivity within the MBON network itself and indirect, convergent feedback from MBONs onto MBINs as possible mechanisms of multimodal sensory integration (*16*), memory consolidation (*17*), and predictive learning (*18*). However, the functional significance of the complex interhemispheric connectivity in this circuit has received less attention. In this study, we take advantage of the anatomical simplicity and genetic accessibility of the larval *Drosophila* olfactory system to ask a fundamental question: how is the representation of sensory input from the two sides of an animal’s nose distributed between the two sides of the brain? Rather than assuming a single locus of interhemispheric integration, we systematically measure the degree of left–right coupling at successive circuit layers, from sensory input to motor-relevant output. This approach allows us to identify where bilateral information is separated, where it is combined, and where it is behaviorally expressed. Our results indicate that rather than maintaining complete functional unilaterality or bilaterality, different output channels of the larval MB preserve or erase stimulus laterality to varying degrees. Moreover, the preserved laterality information can bias navigational decisions, suggesting that this type of multi-decoder architecture may have functional importance for the way that insects process sensory signals to guide behavior.

## 2 Results

### 2.1 Unilateral odor stimulation reveals minimal functional coupling between the two mushroom bodies

To determine where interhemispheric integration arises in the larval olfactory pathway, we developed a strategy for creating unilateral patterns of sensory input. Although analysis of the L1 larval connectome suggests that feedforward connections into the KC layer are predominantly ipsilateral, KCs nevertheless receive contralateral input from two sources: axo-dendritic synapses from the mPN pathway and abundant axo-axonic feedback from MBINs (Fig. 1A and S1). These anatomical features raise the possibility that the two MBs might be functionally coupled despite receiving largely segregated sensory input.

We used a microfluidic platform for simultaneous odor delivery and calcium imaging previously developed in our lab (Fig. 1C; *20*), combined with pulsed UV laser ablation of the antennal nerve to restrict olfactory input to one side of the animal’s head (Fig. 1D). Calcium imaging was performed immediately after ablation to minimize the possibility of compensatory changes in circuit dynamics.

Across five structurally diverse monomolecular odorants, we observed robust odor-evoked responses in KCs ipsilateral to the intact dorsal organ ganglion (Fig. 1E, G–H) but virtually no activity on the contralateral side (Fig. 1F, G–H). Even 1-pentanol, which uniquely recruits the bilateral uPN 35a input channel (*13, 20*), elicited only weak and inconsistent contralateral KC responses. These data indicate that despite the anatomical potential for bilateral coupling, the left and right KCs operate as functionally independent populations.

### 2.2 Modulatory neurons symmetrize reinforcement signals across hemispheres

The finding that KCs respond ipsilaterally raised an immediate question: if odor representations are lateralized at the MB input layer, how does the system ensure that learned associations generalize across hemispheres? The MBIN ensemble—which conveys reinforcement signals capable of modifying KC-MBON synapses—is a natural candidate for mediating such generalization.

We imaged volumetrically from MBIN axon terminals in the MB lobes, using presynaptically trafficked GCaMP6s driven by *Vmat-T2A-GAL4* and taking advantage of the stereotyped com-partmental organization of the MB to extract activity from 9 of 11 larval MB compartments. In striking contrast to the KC responses, MBIN responses showed a high degree of left–right symmetry to one-sided olfactory stimulation: each compartment responded with comparable amplitude to stimulation of either the ipsilateral or contralateral dorsal organ (Fig. 2A–D). After correction for multiple comparisons, no compartment showed a statistically significant difference between ipsi- and contralateral peak response amplitudes (Fig. 2C).

**Figure 2:**
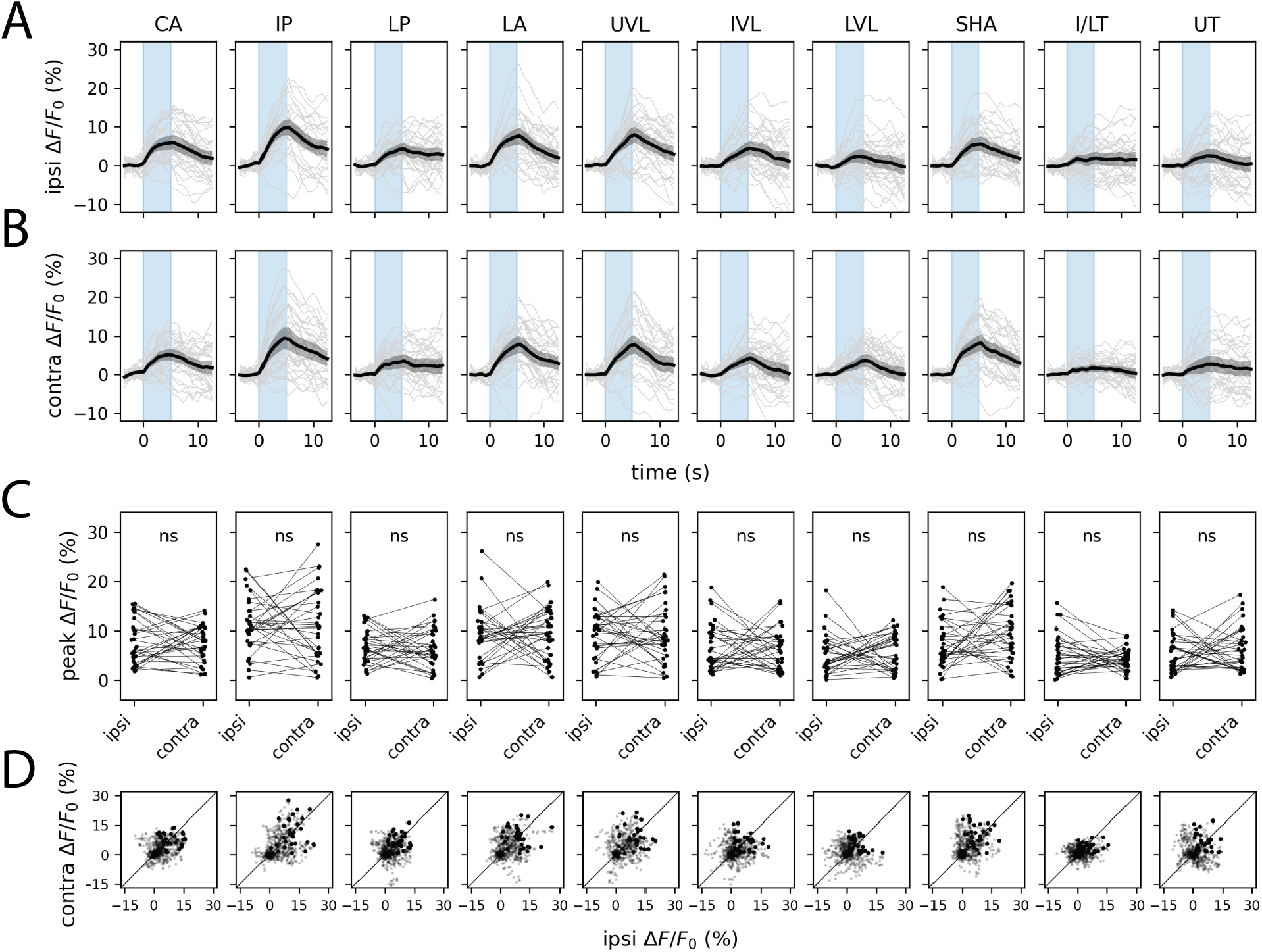
Symmetrization of unilateral olfactory stimuli by the MBIN ensemble. Plots show ipsilateral (A) and contralateral (B) presynaptic GCaMP6s transients and peak ipsilateral vs. contralateral response amplitudes (C) evoked by one-sided stimulation with methyl phenyl sulfide (10^*−*4^). In A and B, dark lines represent grand averages over *n* = 8 animals and 4 stimulus repetitions per animal; gray shaded regions are bootstrapped 95% CIs. Colored vertical bars represent 5 s stimulus periods. D. Scatter plots of ipsilateral and contralateral MBIN activity in the 15 s following stimulus onset. Bold points represent peak responses. Diagonal lines represent equality between ipsilateral and contralateral response amplitudes. Each column shows responses in a different anatomically defined MBIN compartment. One pair of MB compartments, the intermediate toe (IT) and lower toe (LT) of the medial lobe, could not be reliably distinguished and are thus combined for purposes of data visualization. See also Fig. S3. Anatomical abbreviations: CA, calyx; IP, intermediate penduncle; LP, lower peduncle; LA, lateral appendix; UVL, upper vertical lobe; IVL, intermediate vertical lobe; LVL, lower vertical lobe; SHA, shaft; I/LT, intermediate/lower toe; UT, upper toe. Statistics: n.s., *p*_adj_ *≥* 0.05 (paired Student’s *t*-test with Holm’s correction).

### 2.3 MB output neurons exhibit heterogeneous preservation of stimulus laterality

Given that KCs but not MBINs preserve laterality information, we asked what happens at the output layer of the MB. Connectomic analysis predicts considerable heterogeneity: some MBON types project almost exclusively ipsilaterally (MBON-c1) or contralaterally (MBON-m1), while others show mixed innervation patterns (MBON-i1, -a1, -h1/h2) at their dendritic and axonal arbors (Fig. 3A–B; *14, 19, 21*).

**Figure 3:**
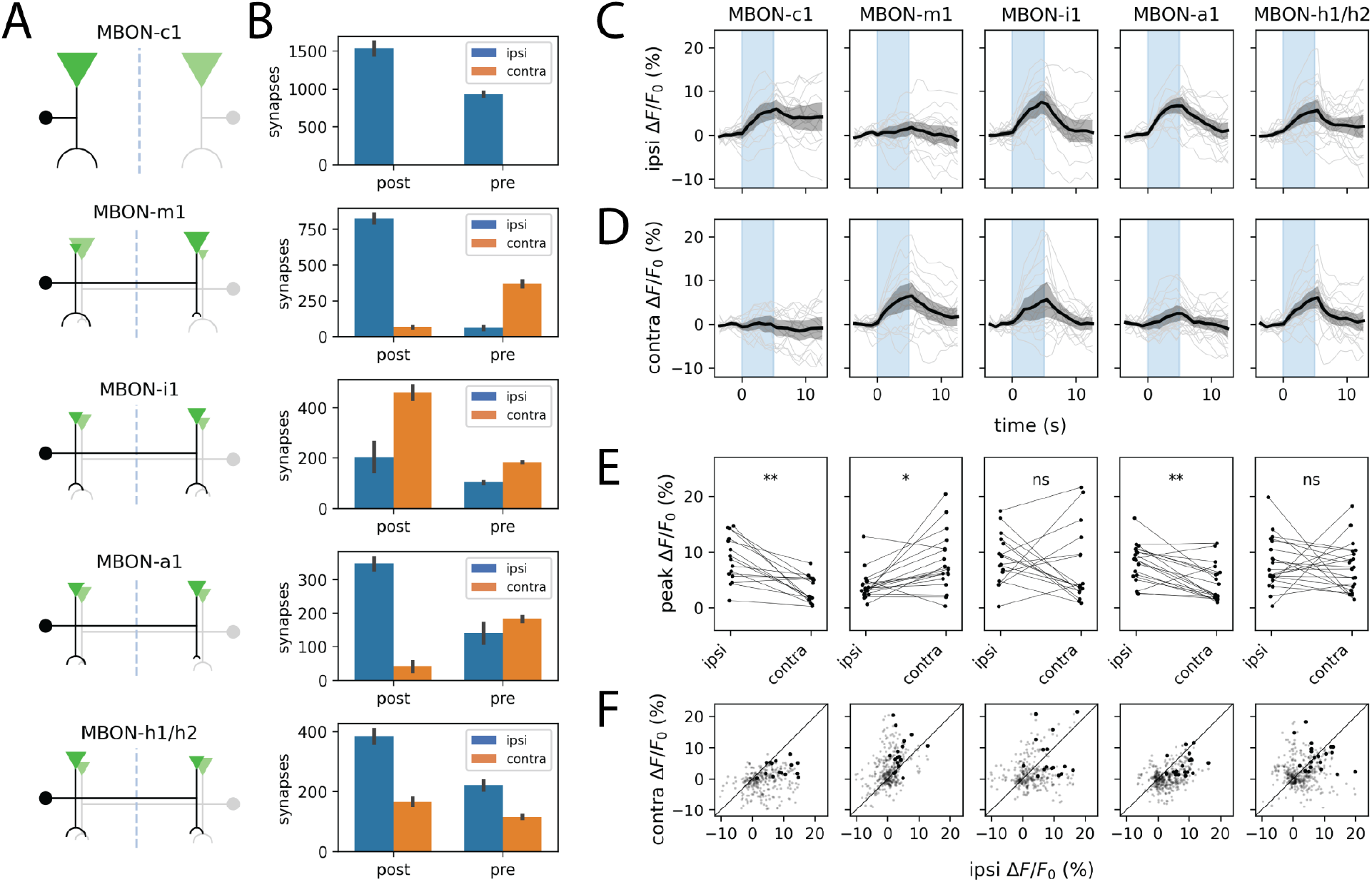
Different MBONs exhibit varying degrees of IHI. A. Schematic projection patterns of 5 different MBON types. Filled circles denote cell bodies. Inverted triangles and semicircular arcs denote pre- and postsynaptic sites, respectively; their size is proportional to the number of such sites per cell type. B. Quantification of ipsi-/contralateral distribution of pre-/postsynaptic sites for each of the MBON types shown in A. Error bars represent s.e.m. (which, for the case of *n* = 2 cells, is simply half of the absolute difference in counts). C–F. MBONs exhibit diverse ipsi-vs contralateral tuning to asymmetric odor stimuli; see also Fig. S4. Plots show ipsilateral (C) and contralateral (D) presynaptic GCaMP6s transients and peak ipsilateral vs. contralateral response amplitudes (E) evoked by one-sided stimulation with methyl phenyl sulfide (10^*−*4^). In C and D, dark lines represent grand averages over *n* = 7 ±1 animals and 3 stimulus repetitions per animal; gray shaded regions are bootstrapped 95% CIs. Colored vertical bars represent 5 s stimulus periods. D. Scatter plots of ipsilateral and contralateral MBIN activity in the 15 s following stimulus onset. Bold points represent peak responses. Diagonal lines represent equality between ipsilateral and contralateral response amplitudes. Each column shows responses in a specific MBON type, labeled by the corresponding split-GAL4 line (see Methods). Statistics: n.s., *p*_adj_ *≥* 0.05; *, *p*_adj_ < 0.05;**, *p*_adj_ < 0.01 (paired Student’s *t*-test with Holm’s correction).

Using split-GAL4 lines targeting five representative MBON subtypes (*22*), we recorded presynaptic calcium responses to unilateral odor stimulation. Consistent with the anatomical predictions, MBON responses ranged from strongly lateralized to nearly symmetric (Fig. 3C–F). MBON-c1 and MBON-m1 almost perfectly preserved information about stimulus side, with minimal signal mixing between hemispheres. MBON-a1 showed a statistically significant ipsilateral bias despite having similar numbers of axon terminals in both hemispheres of the published L1 connectome (*14*). In contrast, MBON-h1/h2 and MBON-i1 exhibited comparable responses to ipsi- and contralateral stimulation.

The strong laterality preservation in MBON-m1 is particularly striking. This neuron has been characterized as a “convergence neuron” rather than a typical MBON, due to the fact that it receives input not only from KCs but also from multiple MBON compartments as well as the LH and is believed to play a role in relaying integrated MB output to premotor areas (*18, 22*). Yet, MBON-m1 faithfully transmits stimulus laterality. This apparent paradox may be resolved by noting that MBON-h1/h2 and MBON-k1—which themselves receive bilateral KC input and make respectively 1- and 2-hop connections onto MBON-m1—likely release the inhibitory neurotransmitters GABA and glutamate (*14, 22*). Thus, the principal *excitatory* inputs onto MBON-m1 are predominantly unilateral.

### 2.4 Lateralized MBON output can bias turning behavior

If some MBONs preserve stimulus laterality five synapses downstream from the sensory periphery, does this information influence behavior? Connectomic analysis reveals that MBONs differ markedly in the laterality of their postsynaptic targets. MBON-m1 is notable for synapsing onto many neurons with strongly asymmetric axon arbors—a wiring motif that potentially supports a role in odor-guided steering (Fig. 4A). We tested this possibility by measuring turn rates during unilateral or bilateral optogenetic activation of individual MBON types. Animals stochastically expressing CsChrimson in neither, one, or both cells of a single left-right MBON pair were stimulated while freely navigating, and their rates of stimulus-triggered ipsilateral and contralateral turns were compared (Fig. 4B–D).

**Figure 4:**
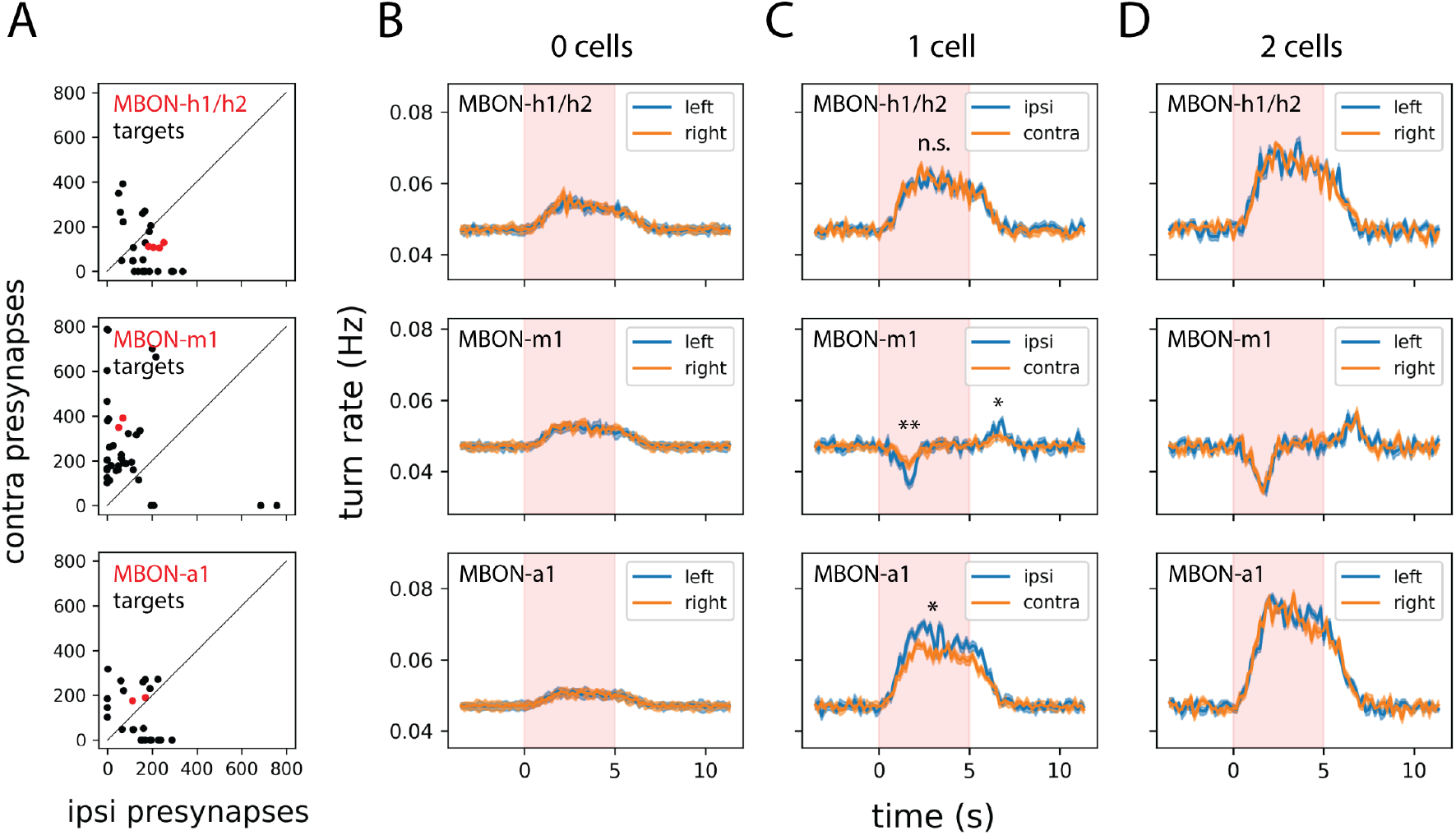
Different MBONs target variously lateralized downstream cell types and exert diverse effects on turning. A. Quantification of the number of ipsi-/contralateral presynaptic sites for all postsynaptic partners of the indicated MBON type. Data for the MBON itself are in red. B–D. Optogenetic activation of individual MBONs modulates turning in a variably side-biased manner; see also Fig. S5 for complete sample size information. Plots depict the change in side-specific turn rates triggered by optogenetic stimulation of 0 (B), 1 (C), or 2 (D) cells of the indicated MBON type in each row. For the 2-cell case of MBON-h1/h2, animals expressed the optogenetic effector CsChrimson::mVenus in 1 cell of each hemisphere. Dark lines represent ensemble averages and shaded regions represent ±1 s.e.m. Statistics: n.s., *p*_adj_ *≥* 0.05; *, *p*_adj_ < 0.05; **, *p*_adj_ < 0.01 (unpaired Student’s *t*-test with Holm’s correction).

Unilateral activation of MBON-m1 or MBON-a1 produced significantly asymmetric effects on turning (Fig. 4C). For the attraction-promoting MBON-m1, activation induced a significantly greater decrease in ipsiversive turning than contraversive turning. For the avoidance-promoting MBON-a1, the pattern was reversed: activation enhanced ipsiversive turning more than contraversive turning. In the case of MBON-m1, an asymmetric OFF-response was also observed: cessation of stimulation induced a significantly greater *increase* in ipsiversive than contraversive turns. These asymmetries were absent in control animals expressing CsChrimson in both hemispheres (Fig. 4B) or neither hemisphere (Fig. 4D). Notably, MBON-h1/h2—which showed symmetric odor responses— produced no turn rate asymmetry upon unilateral activation, consistent with its role in conveying bilaterally integrated rather than lateralized information.

### 2.5 Bilateral spatial comparisons enhance chemotaxis performance

The preservation of laterality information in specific MBONs and its capacity to bias turning suggested that larvae might exploit instantaneous bilateral comparisons during navigation. Although it has been persuasively established that larvae with a single functional ORN can ascend chemical gradients through temporal sampling alone (*23, 24*), this does not preclude an additional role for spatial comparisons when bilateral input is available.

Inspired by previous studies that used optogenetic stimulus gradients to probe the algorithmic basis of larval chemotaxis (*25*), we developed a fictive chemotaxis assay to test this possibility directly. Using stochastic expression of CsChrimson in the attraction-promoting ORNs 42a and 42b, we generated larvae with different numbers and configurations of optogenetically responsive sensory neurons (Fig. 5A–D). Previous work established that these neurons drive quantitatively similar levels of attraction when activated optogenetically (*26*). Of particular interest were larvae with exactly two labeled ORNs, arranged either in a “cis” configuration (both on the same side) or a “trans” configuration (one on each side).

**Figure 5:**
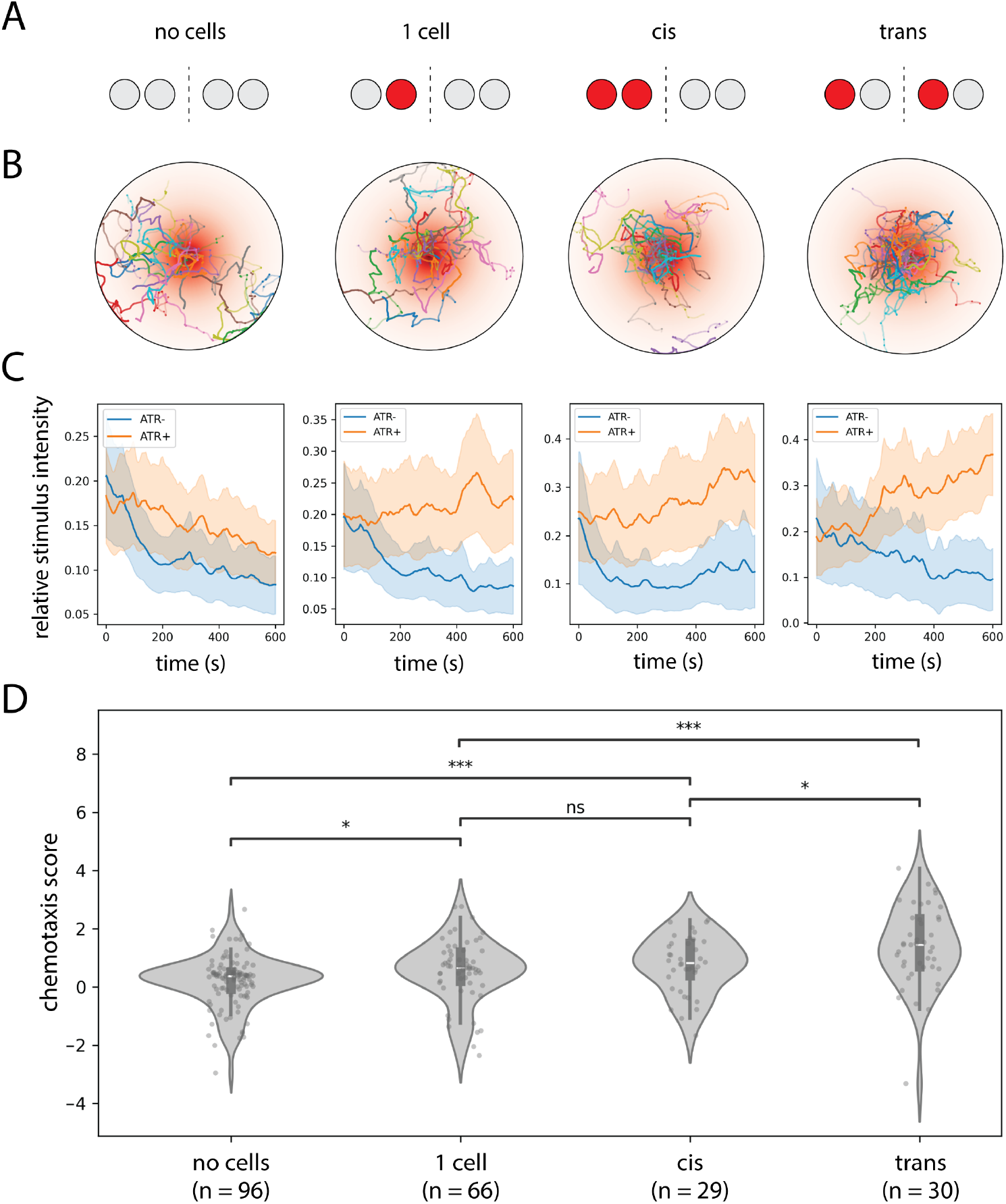
Bilateral spatial comparisons enhance chemotaxis performance. A. Schematic representation of the CsChrimson expression pattern in 42a/42b ORNs for each class of experimental animals. “No cell” larvae express CsChrimson in 0/4 ORNs and “1 cell” larvae in 1/4 ORNs; “cis” and “trans” larvae express CsChrimson in 2/4 ORNs on the same or opposite sides, respectively. B. Spatial trajectories of each class of larvae in a radially symmetric exponential optogenetic gradient.C. Ensemble-averaged time course of fictive stimulus intensity (normalized to maximum value) for each class of larvae. Retinal-fed larvae (ATR+) are shown in orange, control larvae (ATR-) in blue.D. Quantification of chemotaxis scores for each class of larvae. Scores represent the increase in relative stimulus intensity over time for each ATR+ larva normalized to ATR-controls of the same class (see Methods). Larvae in the trans configuration achieve significantly higher mean chemotaxis scores than those in the cis configuration. Statistics: n.s., *p*_adj_ ≥ 0.05; *, *p*_adj_ < 0.05; **, *p*_adj_ < 0.01;***, *p*_adj_ < 0.001 (unpaired Student’s *t*-test with Holm’s correction).

When navigating a radially symmetric exponential light gradient, both configurations successfully ascended toward the stimulus maximum. However, larvae in the trans configuration achieved modestly but significantly higher chemotaxis scores than those in the cis configuration (Fig. 5E;nΔ[chemotaxis score] = 0.30 ± 0.11). In contrast, larvae of the cis configuration were statisticallyn indistinguishable from larvae with only 1 ORN labeled (Fig. 5B). The difference in chemotaxis score between the cis and trans configurations was modest in our experimental conditions but is evidence that larvae can exploit bilateral information provided by both left and right dorsal organs to significantly enhance navigational performance.

## 3 Discussion

Our findings reveal that the larval *Drosophila* olfactory system neither fully integrates nor completely segregates bilateral sensory information. Instead, the MB implements a selective pooling architecture in which some output channels preserve stimulus laterality while others erase it. This design appears to balance two competing computational demands: the need for interhemispheric integration and the potential utility of segregated processing for spatial contrast detection or computational robustness.

By systematically probing successive layers of the olfactory pathway, we identified the MB as the first major site where olfactory signals from the left and right dorsal organs are combined. The KC layer, despite receiving some anatomical input from contralateral sources, operates in a predominantly ipsilateral mode—a finding that underscores the functional independence of the two MBs as associative memory systems. This independence persists surprisingly deeply into the circuit: MBON-m1, positioned five synapses from sensory input and downstream of substantial interhemispheric connectivity, still faithfully reports which side of the head detected an odor.

The heterogeneity we observe in MBON laterality tuning is not simply inherited from the KC layer but is instead imposed by the specific projection patterns of individual MBONs. For example, bilateral MBONs like MBON-h1/h2, which could potentially symmetrize signals onto their post-synaptic targets, instead exert predominantly inhibitory influence—potentially sharpening rather than suppressing left-right contrasts. This arrangement allows convergence neurons like MBON-m1 to integrate multimodal and multi-compartment information while retaining access to the laterality of the original sensory input.

The uniformly symmetric responses of MBINs present a striking contrast to the heterogeneity observed at the MBON layer. We propose that this symmetry reflects a functional requirement for coherent learning: if an odor-punishment association is formed following stimulation of one dorsal organ, the larva should avoid that odor regardless of which side subsequently encounters it. By broadcasting reinforcement signals bilaterally, MBINs may ensure that synaptic plasticity occurs symmetrically across the two MBs, enabling learned associations to generalize across hemispheres.

This interpretation aligns with the known role of MBINs in conveying reward and punishment signals that modulate the strength of KC–MBON synapses (*15*). If MBINs responded asymmetrically to lateralized odor input, the resulting plasticity would be confined to one MB, potentially producing maladaptive hemisphere-specific memory traces. The symmetric architecture we observe may thus represent a solution to the “binding problem” of associating a unilaterally detected odor with a bilaterally experienced reinforcement.

Our optogenetic experiments demonstrate that the laterality preserved in certain MBONs augments the sensory information that is used to effect navigational decision-making. The finding that unilateral activation of MBON-m1 or MBON-a1 biases turning suggests that these neurons may participate in a push-pull steering circuit, consistent with their downstream connectivity to asymmetrically projecting premotor neurons. The fact that the behavioral effect of unilateral activation of these neurons is not perfectly side-specific but merely biased toward one side may reflect residual interhemispheric integration downstream of these circuit nodes. Alternatively, chemical or electrical coupling between the left and right members of each neuron pair could lead to some level of indirect activation of the unlabeled cell.

How laterally-asymmetric MBON activity is used to create a laterally-asymmetric behavioral response natural behavior remains an open question. Fictive chemotaxis in a spatial gradient of optogenetic stimulation is modestly but significantly more effective with two left/right sensory neurons than with two neurons on the same side. If the larva makes instantaneous left/right comparisons of sensory input to improve spatial navigation (i.e., tropotaxis), the laterality preserving MBON channels represent a plausible neural substrate for implementing left/right steering decisions. We note that only a small improvement in navigational is to be expected, as the maximal difference in stimulus intensity between the left and right DOs is quite modest under our experimental conditions. Here, tropotaxis presumably supplements rather than obviates navigational strategies based on temporal integration (klinotaxis).

Nevertheless, the observation that bilateral spatial comparisons can enhance chemotaxis performance adds an additional element to the prevailing model of larval olfactory navigation. Louis *et al*. (*24*) elegantly demonstrated that single-ORN larvae can navigate chemical gradients, leading to the current consensus that temporal sampling of concentration changes during periods of forward movement and during lateral head sweeps constitutes the dominant strategy (*27, 28*). Our results do not contradict this view but suggest that when bilateral input is available, it provides additional information that larvae can exploit. Interestingly, larval phototaxis is a case of navigational behavior driven by strategies that exploit both tropotaxis and klinotaxis (*29*).

The relatively small magnitude of the trans vs. cis advantage in our assay may reflect the shallow gradients we employed, chosen to resemble smooth spatial gradients used in previous studies of airborne odor navigation. For larvae directly immersed in rotting fruit or other semi-liquid substrates and sensing dissolved rather than airborne odorants, olfaction may arguably function more like a contact sense, conceivably leading to much steeper stimulus gradients than those tested here. Even so, the adult fly is known to be able to detect instantaneous gradients in airborne odor intensity and even infer the direction of odor motion from the differential signal at its two antennae, which have a spatial separation only twice that of the dorsal organs of an L3 larva (*12, 30, 31*).

Several limitations of our study deserve mention. First, it is possible that the dynamics of neural activity in the larval olfactory system are somewhat different in freely moving compared to immobilized animals. Although the conclusions from our calcium imaging experiments comport with the results of our behavioral and connectomic analyses, we cannot rule out that response properties of neurons in the larval olfactory system might be influenced by internal state signals accompanying passive immobility. Our behavioral assays also deviate from fully lifelike conditions in their use of highly simplified stimulus geometries which may fail to capture important aspects of real odor stimuli in the animal’s natural habitat. Indeed, little is known about the statistical structure of naturalistic odor landscapes on the physical scales relevant to fly larvae. Finally, our imaging experiments were conducted exclusively in L1 larvae, and it remains to be determined how the functional architecture of left/right integration is maintained or reorganized during subsequent larval stages.

Even so, the selective interhemispheric architecture we describe in the larval MB may reflect a general solution to a problem faced by many bilateral sensory systems: how to integrate information for unified perception while retaining access to spatial structure for oriented behavior. In vertebrate auditory systems, sound localization depends on circuits that compute interaural timing and intensity differences, while higher-order areas construct unified auditory objects (*3*). In mammalian olfaction, recent work has revealed that piriform cortex contains both bilateral and unilateral neurons, with the balance potentially shaped by experience (*11*). The circuit mechanisms we have uncovered in the larval olfactory system may therefore exemplify fundamental principles governing how bilaterian nervous systems balance the competing demands of unified perception and spatial computation.

## 4 Materials and methods

### 4.1 Fly husbandry and genetics

Fruit flies were maintained at 22 ^*°*^C with 60% relative humidity and a 12/12 light/dark cycle in vials with standard cornmeal-based food. To collect larvae, we allowed adult flies to lay eggs on grape juice agar plates with fresh yeast paste, following established protocols (*32*). We conducted calcium imaging using late L1 larvae (2 days after egg laying [AEL]), while behavioral experiments used early L3 larvae (3–4 days AEL). Our imaging studies utilized two genetic effector lines: *UAS-GCaMP6m*; *Orco-RFP* (Fig. 1) and *UAS-syt::GCaMP6s, MB247-RFP*; *Orco-RFP* (for all other experiments). The following driver lines were used to target genetically defined subsets of neurons:

| cell type | driver line |
| --- | --- |
| KCs | *R14H06-GAL4* |
| MBINs | *Vmat-T2A-GAL4* |
| MBON-a1 | *SS02006 split-GAL4* |
| MBON-c1 | *SS01776 split-GAL4* |
| MBON-h1/h2 | *SS00894 split-GAL4* |
| MBON-i1 | *SS01726 split-GAL4* |
| MBON-m1 | *SS02163 split-GAL4* |
| ORN 42a | *Or42a-GAL4* (III) |
| ORN 42b | *Or42b-GAL4* (II) |

For behavioral experiments involving optogenetic activation of individual MBONs (4), we used a stochastic labeling approach to produce animals expressing an mVenus-tagged CsChrimson transgene in a subset of cells labeled by a given MBON split-GAL4 line:

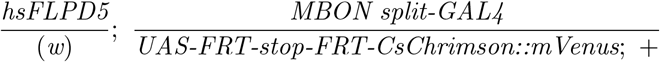

Animals of this genotype were behaviorally profiled (see below) and subsequently examined under an inverted confocal microscope to identify the number of MBONs labeled (0, 1, or 2). Animals were raised at 25 ^*°*^C to induce expression of FLP recombinase.

An analogous strategy was used for behavioral experiments involving stochastic optogenetic activation of ORNs (5), involving animals of the genotype:

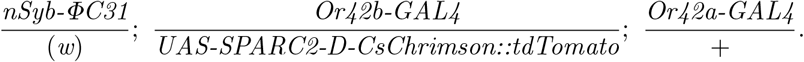

For this experiment, the use of SPARC allowed more stable control over the proportion of animals expressing CsChrimson in different subsets of the Or42a/Or42b ORNs.

### 4.2 Functional imaging and microfluidics

For simultaneous odor delivery and calcium imaging of intact first-instar larvae, we employed a previously documented microfluidic system (*33*). All experiments were conducted using an 8-channel microfluidic chip. The experimental protocol involved 5 s odor exposures interleaved with 30 s water wash-out periods. We selected appropriately sized larvae directly from grape plates, washed them three times with distilled water, and inserted them into the microfluidic device using a 1 mL syringe containing 0.1% Triton X-100 solution. Each larva was carefully positioned with its dorsal side facing the microscope lens at the end of a narrow channel. We captured fluorescent signals using an EMCCD camera (Andor iXon) connected to a Yokogawa inverted spinning-disk confocal microscope (Nikon). All live imaging was performed with a 40X/1.15 water immersion objective (Nikon). We used a piezo scanner (PIFOC) to acquire 3D volumes consisting of approximately 18 ± 4 *z*-planes spaced 1.5 µm apart, at a volume scanning rate of (0.9 ± 0.2) Hz.

### 4.3 Laser ablation

A 365 nm pulsed dye laser (Andor Micropoint) tuned to 435 nm and integrated into the light path of our microscope was used to ablate the antennal nerve near its exit from the dorsal organ ganglion (DOG). Trains of 10 pulses (each approximately 170 µJ) were delivered at 10 Hz until the nerve visibly ruptured. The ORN axons were genetically labeled via *Orco-RFP* for ease of targeting. The side ablated (left or right) was randomized across animals, allowing ipsilateral and contralateral data to be pooled across left/right experiments. Calcium imaging experiments were performed immediately after ablation to minimize the possibility of homeostatic compensatory changes following this perturbation.

### 4.4 Behavior and optogenetics

Single-animal optogenetic assays were performed by placing larvae in individual agarose-coated arenas of a 24-arena assay plate. This plate was then inserted into a behavior rig equipped with an overhead projector and IR camera, under the control of a custom MATLAB script. Light from the projector was passed through a 600 nm filter (Edmund Optics) to minimize visual stimulation. Larval trajectories were tracked via the MARGO software package (*34*) and analyzed with MA-GATAnalyzer (*28*). For these experiments, larvae were reared on yeast paste supplemented with all-trans retinal (100 mM). The illumination intensity was 5 µW mm^*−*2^. For fictive tropotaxis experiments (Fig. 5), the pattern of illumination was a radially symmetric exponential gradient (length scale 8 mm) of peak intensity 5 µW mm^*−*2^ at the center of each arena. In Fig. 5, the chemotaxis score for larva *i* was calculated according to the formula

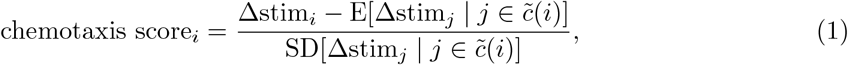

where Δstim_*i*_ is the net change in the fictive stimulus level experienced by larva *i* over the course of the assay, and 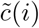 represents the set of ATR^*−*^ larvae in the same expression class as larva *i*.

### 4.5 Image analysis

To process volumetric calcium recordings, we first corrected for motion using groupwise nonlinear registration with the `itk-elastix` software package (*35*). We then manually annotated regions of interest (ROIs) for the target cells and/or MB compartments in the time-averaged 4D volumes (three spatial dimensions plus color channel) using `napari` and custom python code (*36*). We extracted time series data by taking the 90th percentile of raw fluorescence signal from each ROI and calculated Δ*F/F*_0_ traces. The baseline fluorescence (*F*_0_) was determined by averaging the signal during the 5 seconds immediately before stimulus onset, while peak fluorescence was measured as the maximum value of *F* during the stimulus period.

## 5 Acknowledgments

We thank current and former members of the Samuel and de Bivort labs for valuable assistance and helpful discussions, especially K. Vogt, G. Si, J.K. Kanwal, S. Lazopulo, H. Casademunt, I.S. Chandok, N. Rodman, and M. Kim. We are also grateful to M. Zlatic and N. Hu for sharing transgenic fly strains. Stocks obtained from the Bloomington *Drosophila* Stock Center (NIH P40OD018537) were used in this study. Microfluidic devices were fabricated at the Harvard University Center for Nanoscale Systems; a member of the National Nanotechnology Coordinated Infrastructure Network (NNCI), which is supported by the NSF (ECCS-2025158).

## 5.2 Author contributions

Conceptualization: D.M.Z., B.L.d.B., and A.D.T.S. Methodology: D.M.Z. Software: D.M.Z. Validation: D.M.Z. Formal analysis: D.M.Z. Investigation: D.M.Z., B.L.d.B., and A.D.T.S. Data curation: D.M.Z. Resources: D.M.Z. Writing—original draft: D.M.Z. Writing—review and editing: D.M.Z., B.L.d.B., and A.D.T.S. Visualization: D.M.Z. Supervision: B.L.d.B. and A.D.T.S. Project administration: A.D.T.S. Funding acquisition: A.D.T.S.

## 5.4 Data and materials availability

All data needed to evaluate the conclusions in the paper are present in the paper and/or the Supplementary Materials.

