## Supplemental figures for "The larval *Drosophila* mushroom body balances lateralized sensing and interhemispheric integration"

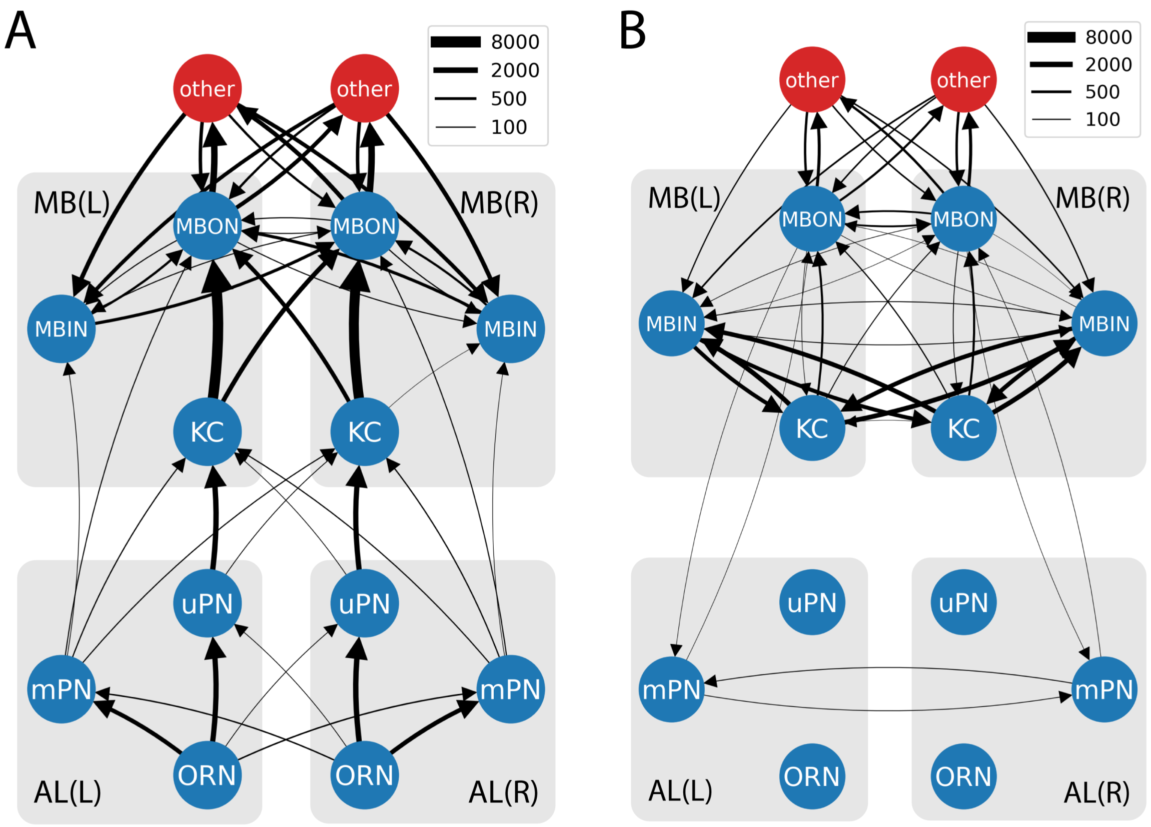


**Figure S1: Axo-dendritic and axo-axonic connectivity of the larval olfactory system.** Summarized wiring diagrams of the L1 larval olfactory system, analogous to Fig. 1A except that only axo-dendritic (A) or axo-axonic (B) connectivity is shown. Data from Winding *et al.* (*1*). Related to Fig. 1.

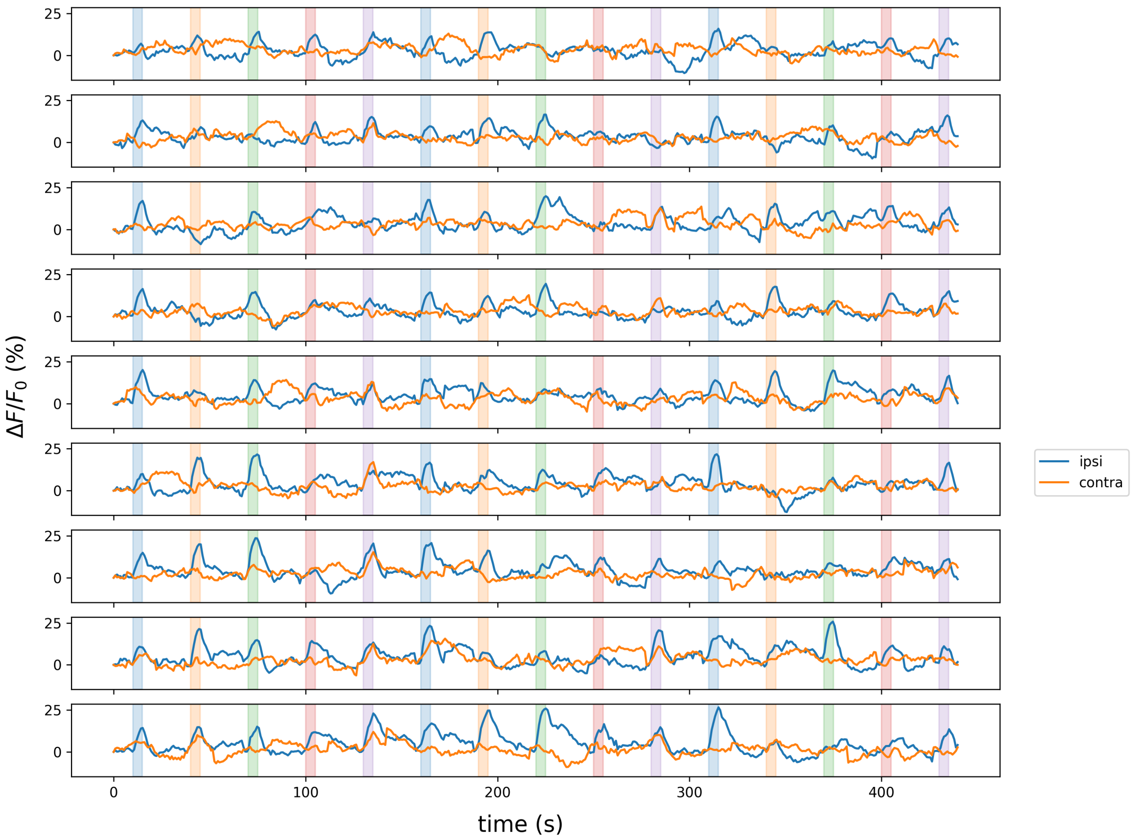


**Figure S2: Raw calcium responses of ipsi- and contralateral KCs to unilateral olfactory stimulation.** Full calcium response traces for each animal from Fig. 1E–H. Each row represents a different larva ($n = 3$ technical replicates from each of 9 animals). The side ablated (left or right) was randomized across animals. Shaded bars indicate odor periods. Color code: MePhS, methyl phenyl sulfide (blue); EtOAc, ethyl acetate (orange); BzCHO, benzaldehyde (green); GeOAc, geranyl acetate (red); 1PnOH, 1-pentanol (violet). All odors were used at $10^{-4}$ dilution. Related to Fig. 1.

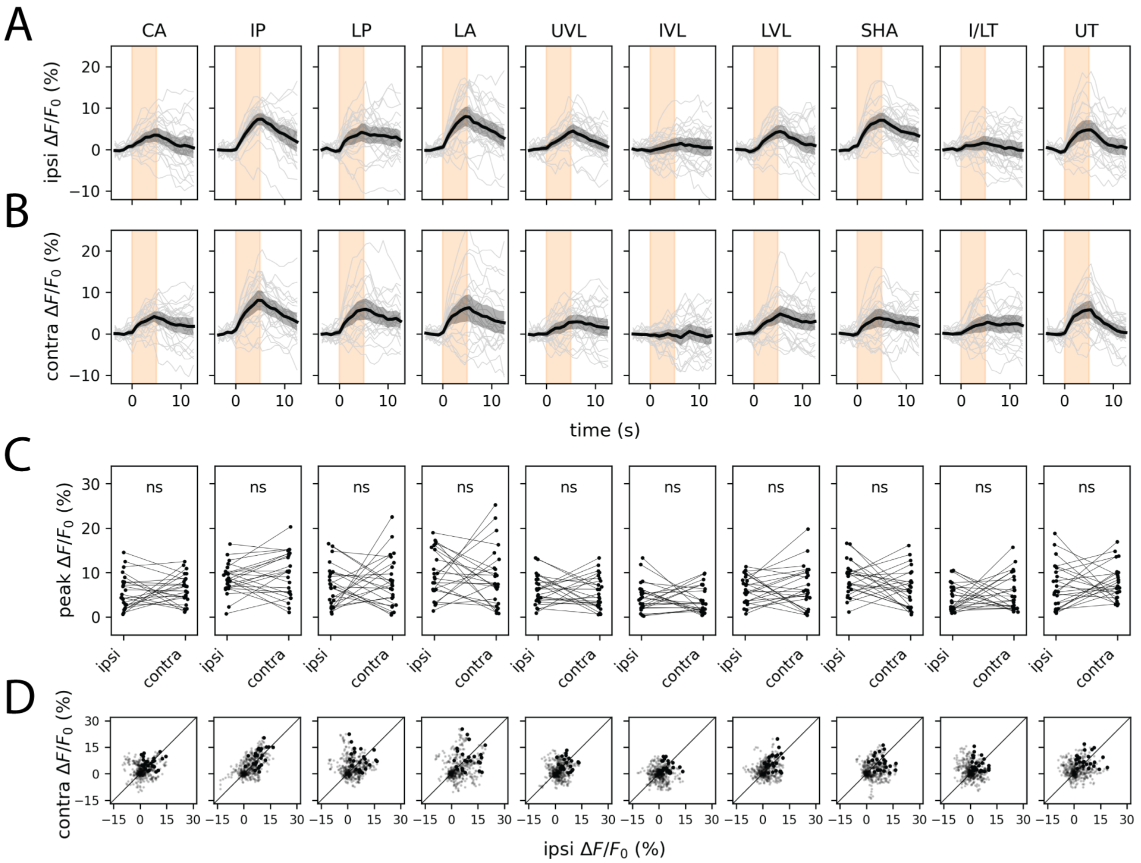


**Figure S3: Symmetrization of unilateral olfactory stimuli by the MBIN ensemble (alternate odorant).** Plots show ipsilateral (A) and contralateral (B) presynaptic GCaMP6s transients and peak ipsilateral vs. contralateral response amplitudes (C) evoked by one-sided stimulation with ethyl acetate ($10^{-4}$). In A and B, dark lines represent grand averages over $n = 8$ animals and 4 stimulus repetitions per animal; gray shaded regions are bootstrapped 95% CIs. Colored vertical bars represent 5 s stimulus periods. D. Scatter plots of ipsilateral and contralateral MBIN activity in the 15 s following stimulus onset. Bold points represent peak responses. Anatomical abbreviations: CA, calyx; IP, intermediate penduncle; LP, lower peduncle; LA, lateral appendix; UVL, upper vertical lobe; IVL, intermediate vertical lobe; LVL, lower vertical lobe; SHA, shaft; I/LT, intermediate/lower toe; UT, upper toe. Statistics: n.s., $p_{\text{adj}} \geq 0.05$ (paired Student's $t$-test with Holm's correction). Related to Fig. 2.

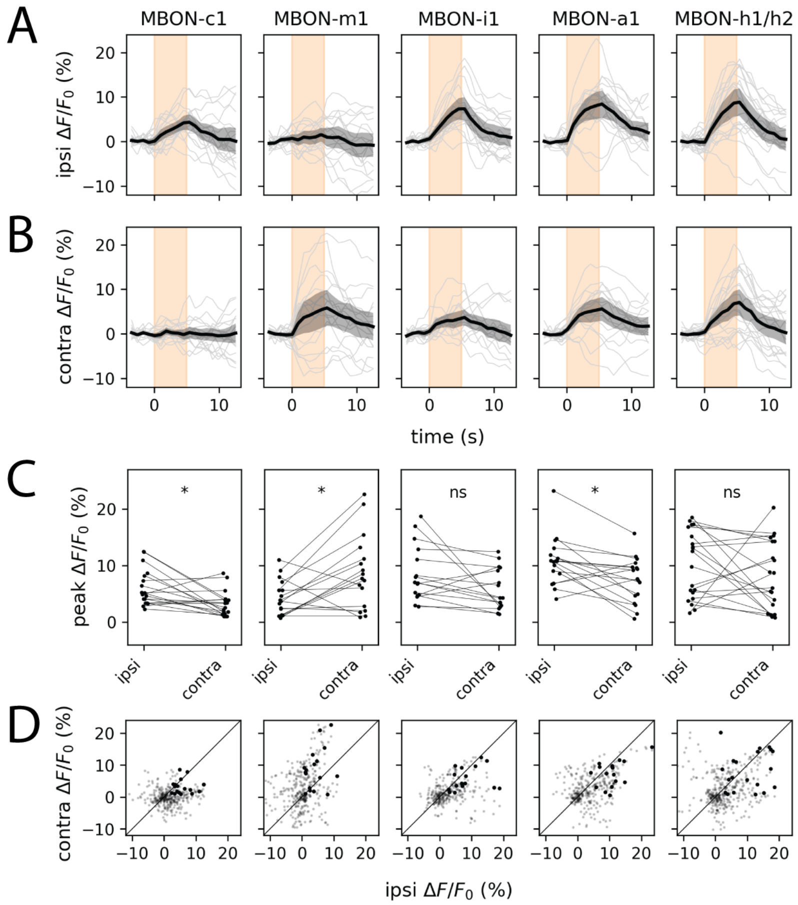


**Figure S4: MBONs exhibit diverse ipsi- vs contralateral tuning to asymmetric odor stimuli (alternate odorant).** Plots show ipsilateral (A) and contralateral (B) presynaptic GCaMP6s transients and peak ipsilateral vs. contralateral response amplitudes (C) evoked by one-sided stimulation with ethyl acetate ($10^{-4}$). In A and B, dark lines represent grand averages over $n = 7 \pm 1$ animals and 3 stimulus repetitions per animal; gray shaded regions are bootstrapped 95% CIs. Colored vertical bars represent 5 s stimulus periods. D. Scatter plots of ipsilateral and contralateral MBIN activity in the 15 s following stimulus onset. Bold points represent peak responses. Diagonal lines represent equality between ipsilateral and contralateral response amplitudes. Each column shows responses in a specific MBON type, labeled by the corresponding split-GAL4 line (see Methods). Statistics: n.s., $p_{\mathrm{adj}} \geq 0.05$; *, $p_{\mathrm{adj}} < 0.05$; **, $p_{\mathrm{adj}} < 0.01$ (paired Student's $t$-test with Holm's correction). Related to Fig. 3.

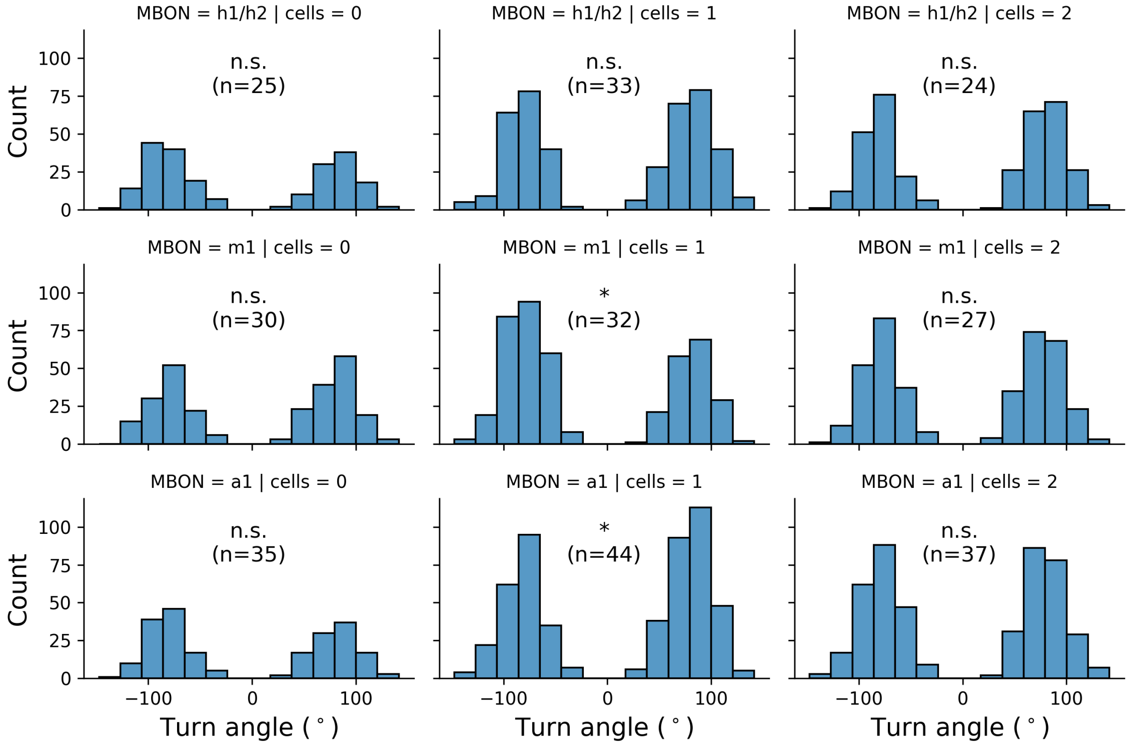


**Figure S5: Optogenetic activation of individual MBONs modulates turning in a variably side-biased manner.** Histograms of turn angle (pooled over all detected turns by all animals within each class) triggered by optogenetic stimulation of 0, 1, or 2 cells (columns) of the indicated MBON type (rows). For the 2-cell case of MBON-h1/h2, animals expressed the optogenetic effector CsChrimson::mVenus in 1 cell of each hemisphere. Statistics: n.s., $p_{\text{adj}} \geq 0.05$; *, $p_{\text{adj}} < 0.05$ (Wilcoxon signed-rank test with Bonferroni correction). Related to Fig. 4.
